# An Alternatively Spliced Gain-of-Function NT5C2 Isoform Contributes to Chemoresistance in Acute Lymphoblastic Leukemia

**DOI:** 10.1101/2023.09.14.557413

**Authors:** Manuel Torres-Diz, Clara Reglero, Catherine D Falkenstein, Annette Castro, Katharina E. Hayer, Caleb M. Radens, Mathieu Quesnel-Vallières, Zhiwei Ang, Priyanka Sehgal, Marilyn M. Li, Yoseph Barash, Sarah K. Tasian, Adolfo Ferrando, Andrei Thomas-Tikhonenko

## Abstract

Relapsed or refractory B-cell acute lymphoblastic leukemia (B-ALL) is a major cause of pediatric cancer-related deaths. Relapse-specific mutations do not account for all chemotherapy failures in B- ALL patients, suggesting additional mechanisms of resistance. By mining RNA-seq datasets of paired diagnostic/relapse pediatric B-ALL samples, we discovered pervasive alternative splicing (AS) patterns linked to relapse and affecting drivers of resistance to glucocorticoids, anti-folates, and thiopurines. Most splicing variations represented cassette exon skipping, “poison” exon inclusion, and intron retention, phenocopying well-documented loss-of-function mutations. In contrast, relapse-associated AS of NT5C2 mRNA yielded an isoform with the functionally uncharacterized in-frame exon 6a. Incorporation of the 8-amino acid sequence SQVAVQKR into this enzyme created a putative phosphorylation site and resulted in elevated nucleosidase activity, which is a known consequence of gain-of-function mutations in NT5C2 and a common determinant of 6-mercaptopurine (6-MP) resistance. Consistent with this finding, NT5C2ex6a and the R238W hotspot variant conferred comparable levels of resistance to 6-MP in B-ALL cells both in vitro and in vivo. Furthermore, both the NT5C2ex6a and R238W variants induced collateral sensitivity to the inosine monophosphate dehydrogenase (IMPDH) inhibitor mizoribine. These results ascribe an important role for splicing perturbations in chemotherapy resistance in relapsed B-ALL and suggest that IMPDH inhibitors, including the commonly used immunosuppressive agent mycophenolate mofetil, could be a valuable therapeutic option for treating thiopurine-resistant leukemias.

## Introduction

Leukemia is the most common cancer in children and adolescents/young adults, accounting for almost 1/3 of all pediatric cancers, with the majority of cases classified as B-cell acute lymphoblastic leukemia (B-ALL). Only ∼50% of children with first relapse survive long term, and outcomes are appreciably poorer with subsequent relapses. Early bone marrow relapses occurring within three years of diagnosis are particularly difficult to salvage (1). As a result, relapsed or refractory (r/r) high-risk B-ALL cases account for a substantial number of pediatric cancer-related deaths. In addition, while most cases of B- ALL are diagnosed early in life, most deaths from B-ALL (∼80%) occur in adults, largely due to r/r disease. Adults with B-ALL experience very high relapse rates and long-term event-free survival of less than 50% (2,3).

What molecular mechanisms are responsible for treatment failures in r/r B-ALL is only partially understood. Previous whole exome/genome sequencing efforts have identified a dozen of recurrent mutations in relapse-specific targets (4-6). While some of them (e.g., *TP53*) are known to mediate responses to pan-cancer drugs such as anthracyclines, others are involved in therapeutic responses to specific leukemia therapeutics: glucocorticoids (*NR3C1, WHSC1*), anti-folates (*FPGS*), and thiopurines (*PRPS1, PRPS2, NT5C2*). The most recent effort to catalog such mutations yielded up to 50 genes significantly enriched at first relapse but often as subclonal or low-recurrence events (7). Even the most recurrently affected drivers are mutated in fewer than 20% of all samples suggesting a major role for an alternative mechanism of gene dysregulation. One such mechanism is alternative spicing (AS); yet its role in acquired resistance to chemotherapy remains poorly defined.

To bridge this gap, we previously had generated and mined new and existing RNA-seq datasets in search for local splicing variations prevalent in B-ALL samples, but rare in normal bone marrow cell counterparts. Of note, B-ALL-specific AS was found to affect 15 out of the 20 top leukemia driver genes from the COSMIC database with frequencies far exceeding those of somatic mutations (8). This intersection suggests that AS could be an important mechanism of both leukemogenesis and drug resistance. Here we provide evidence that a subset of relapses often lacking identifiable acquired mutations has an AS signature simultaneously affecting multiple r/r B-ALL drivers. Among them is a novel NT5C2 exon 6a mRNA variant encoding a gain-of-function nucleotidase isoform with increased enzymatic activity capable of conferring resistance to 6-mercaptopurine (6-MP).

## Materials and Methods

### Splicing analysis

The Modeling Alternative Junction Inclusion Quantification (MAJIQ) algorithm version 4.4 [(9); (RRID:SCR_016706)] was run with each sample being quantified individually on a build comprising 219 TARGET B-ALL samples with *de novo* junction detection, but without intron retention detection, as described previously (10-13). Samples were run against the gencode hg38 quantification. For ΔPSI (percent-spliced-in) quantification, the MAJIQ PSI and Voila tools were used on 48 diagnostic-relapse paired samples to extract all the LSVs in all the samples with the –show-all flag. Visualization and downstream analyses were conducted in R using the ggplot2 (RRID:SCR_014601), ComplexHeatmap (RRID:SCR_017270), and tidyverse (RRID:SCR_019186) packages. The sashimi plots were generated with the ggsashimi tool. In parallel, raw junction-spanning reads were obtained from the STAR aligner (see Supplementary Methods) and expressed as JPMs (junction counts per million). JPMs for select AS events in select samples are included in Supplementary Table S1.

### Oxford Nanopore Technologies (ONT) targeted long-read sequencing

For each target gene, primers corresponding to first and last exons were designed to ensure full coverage of relevant alternative splicing events. Typically, 5ng of cDNA were amplified with the long LongAmp Taq 2X Master Mix (New England Biolabs) for 25 cycles. The resulting amplicons were subjected to amplicon-seq (#SQK-NBD114.24, ONT) library preparation. Subsequently, each library was loaded into a Spot- ON flow cell R10 Version (#FLO-MIN114, ONT) and sequenced in a MinION Mk1C device (ONT) until at least 1000 reads per sample were obtained. Results were aligned using Minimap2 version 2.24- r1122 (RRID:SCR_018550) and visualized in Integrative Genomics Viewer (IGV; RRID:SCR_011793) version 2.12.3. The raw sequencing data are publicly available in the Sequence Read Archive (SRA) as Project PRJNA1138448: https://www.ncbi.nlm.nih.gov/sra/PRJNA1138448.

### Cell lines and cell culture

REH (RRID:CVCL_1650), Nalm-6 (RRID:CVCL_0092), 697 (RRID:CVCL_0079), and MHH-CALL-4 (RRID:CVCL_1410) human B-ALL cell lines originally obtained from commercial repositories were cultured and maintained in conditions recommended by ATCC or DSMZ. Cell line authentication for REH and MHH-CALL-4 was performed by short tandem repeat profiling. All cell lines were routinely subjected to Sartorius EZ-PCR™ Mycoplasma detection kit testing.

### Genome editing

sgRNAs targeting CREBBP exons 25 and 26 and CAS9 protein were obtained from IDT (see Supplementary Methods). Cas9 ribonucleoprotein (RNP) complexes were assembled following manufacturing recommendations. These RNP were transfected into 697 and REH cells via electroporation using the Neon transfection system (Invitrogen Neon Transfection System Model MPK5000) and the Neon 10uL Transfection kit (#MPK1096 Invitrogen) using the following conditions: 1700V, 20ms, 2 pulses. Their effects on exon removal were measured by cDNA PCR with primers amplifying the region between exons 20 and 27 (see Supplementary Methods) and Western blotting.

### Morpholino treatment

For FPGS splice switching a specific Vivo-Morpholino targeting the canonical 3’ splice site of Exon 8 was designed (AAATTCCACTGGTCCGTCTGACCCC). As a control, the reverse sequence was used in order to maintain sequence composition, as recommended by the manufacturer. Cells were incubated with the 2.5 μM morpholinos for 24h before treatment with MTX.

### Plasmid constructs and viral infections

For NT5C2 constructs, the coding sequence of the canonical NT5C2 isoform (NM_001351169.2) was amplified using specific primers (see Supplementary Methods) and cloned into the pLVX backbone (Clontech) modified with the EF1a promoter and the Basticidin resistance gene using NEBuilder HiFi DNA Assembly Master Mix (New England Biolabs). Specific primers and NEBuilder HiFi DNA Assembly Master Mix were used to introduce exon 6a or the R238W mutation into the canonical sequence of NT5C2. For viral particles production, 293T cells (ATCC) grown to 90% confluence in a 10cm Petri dish were transfected with 10μg of the NT5C2 constructs, 2.5μg of the pMD2.G (#12259 Addgene), and 7.5μg of the psPAX2 (#12260 Addgene) using 60μL of 25μM PEI (#24765 Polysciences). Supernatants were collected 48h after transfection, passed through 0.45μm PVDF filters, and incubated for 48h with 4×10^6^ cells in media containing 10µg/mL plolybrene (#HY-112735 MedChemExpress). After 48h, culture media were replaced, and cells were incubated with 10μg/mL blasticidin for 96h.

### NT5C2 recombinant protein production and purification

Recombinant wild-type and mutant NT5C2 constructs were cloned, expressed, and purified as previously described (14). Briefly, the full-length NT5C2 cDNA constructs with N-terminal hexahistidine (His6) tags were inseted into the pET28a-LIC expression vector. Recombinant proteins were expressed in Rosetta 2 (DE3) E. coli cells by induction with 0.5 mmol/L isopropyl-β-D-thiogalactopyranoside overnight at 16°C. Then recombinant proteins were purified using an ÄKTA fast protein liquid chromatography system (GE Healthcare). This was followed by affinity and size exclusion chromatography.

### 5′-Nucleotidase assays

5′-nucleotidase activities of purified wild-type and mutant NT5C2 proteins were measured in the absence and presence of allosteric activators using the 5′-NT enzymatic test kit (Diazyme) according to the manufacturer’s instructions, as described previously (15). All assays were performed in triplicate in a Glomax multi-detection system plate reader (Promega).

### In vitro killing assay

For methotrexate (#S1210 Selleck Chemicals), 6-mercaptopurine (#852678-5G- A Sigma Aldrich), mizoribine (#S1384 Selleck Chemicals), doxorubicin (#HY-15142 MedChem Express), dexamethasone (#D4902 Sigma Aldrich), and vincristine (#HY-N0488 MedChem Express) treatments, cells were incubated with increasing concentrations of each drug for 72 hours. Vehicle control dilutions corresponding to the highest drug concentration were used as a baseline measure for 100% survival. Cell viability was measured by CellTiter-Glo assays (G7572 Promega) following manufacturer’s instructions. IC_50_ values were calculated using Prism software v9.3.1 (GraphPad software) with the log (inhibitor) vs. normalized response – variable slope.

### In vivo cell line xenograft animal studies

Luciferase-transduced REH cells were transduced with NT5C2 wild-type, E6a, or R238W constructs and selected with blasticidin for 96h. Following selection, NOD/SCID/Il2rgtm1wjl/SzJ (NSG) mice were intravenously injected with 1×10^6^ parental or modified REH cells and assessed for leukemia engraftment using an In Vivo Imaging System (Xenogen). 10 days after B-ALL engraftment was detected, cohorts of mice (n=5/group) of each REH cell type were randomized for treatment with vehicle control or 6-mercaptopurine (100 mg/kg/day). 6-MP was administered intraperitoneally daily x 5 days as described previously (16). Radiance as a surrogate for total leukemia burden was measured by in vivo bioluminescent imaging (BLI) of mice following intraperitoneal injection of luciferin substrate. BLI assessment was performed with Living Image software (Perkin-Elmer) with statistical analysis and data display performed using Prism (GraphPad).

### Data availability

Main RNA-seq datasets analyzed in this study were parts of the TARGET ALL Phase I and II projects (https://portal.gdc.cancer.gov/projects) and the BLUEPRINT consortium (http://dcc.blueprint-epigenome.eu). TARGET datasets accessed through dbGaP were phs000463.v23.p8 Acute Lymphoblastic Leukemia (ALL) Pilot Phase 1 and phs000464.v23.p8 Acute Lymphoblastic Leukemia (ALL) Expansion Phase 2. All other raw data generated in this study are available upon request from the corresponding author.

## Results

### A subset of r/r B-ALL has an AS signature affecting multiple chemoresistance genes

To gain new insights into molecular mechanisms that render B-ALL chemotherapy ineffective, we analyzed 48 TARGET patients with available paired diagnosis-relapse (D-R) RNA-seq datasets. We called mutations in known r/r driver genes from the RNA-seq data using Genome Analysis Toolkit (GATK; see Supplementary Methods). In agreement with prior analyses, we observed that in 12 out of 48 D-R pairs (25%) there were no identifiable relapse-specific mutations in 12 common r/r genes (Figure 1a). To determine whether differential gene expression (DGE) could account for the chemoresistant phenotype, we performed the Principal Component Analysis on all 96 samples. We observed no separation of diagnostic and relapse samples (Supplementary Figure 1a, red and cyan dots, respectively), suggesting that DGE as a whole is not a primary driver of B-ALL relapses. However, pervasive AS reported in our earlier B-ALL study (8) could contribute to chemoresistance by altering expression levels of distinct mRNA isoforms. Thus, we asked whether any members of the HNRNP and SRSF/TRA2 superfamilies of splicing factors (SF) strongly implicated in exon inclusion/skipping are affected by mutations or copy number alterations (CNA). Using cBioPortal (17), we identified such CNAs in 14 out of 48 patients (30%). For example, deep deletions of both SRSF4 and SRSF10 genes was found in patient PAPZST and the SRSF3 gene - in samples PARAKF and PARJSR (Figure 1b). Of note, these deletions affected only the latest among longitudinal samples, attesting to their possible role in B-ALL relapses.

**Figure 1.**
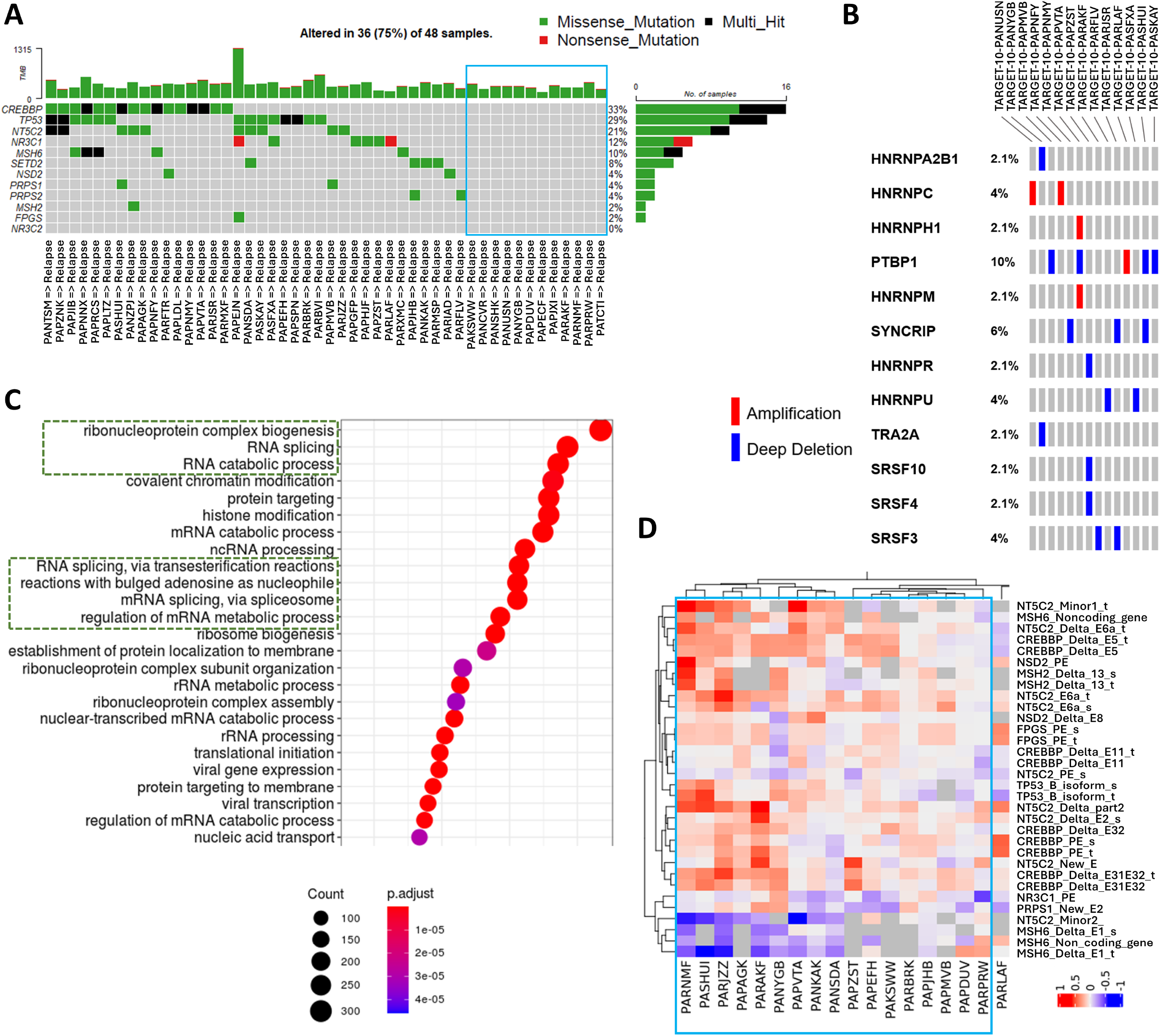
Splicing signature of relapsed leukemias. **a**. Oncoprint showing distribution of acquired mutations in known relapse/resistance genes in 48 paired TARGET samples. The blue rectangle on the right marks samples with no identifiable relapse-specific mutations. **b**, Oncoprint showing distribution of mutations in select splicing factor genes in 48 TARGET patients under investigation. **c**. Gene Set Enrichment Analysis of transcripts comprising one of the unsupervised clusters. Green rectangles denote GO categories involving RNA splicing. **d**. Heatmap showing changes in splicing in a cluster of 17 relapse vs. diagnostic B-ALL samples (light blue box). The dendrograms on top represent the results of unsupervised clustering without excluding samples with known relapse-specific mutations. The adjacent sample PARLAF with several profoundly mis-spliced transcripts is shown for comparison. The red and blue colors represent increases/decreases in inclusion of exonic segments, respectively.

To explore the extent and functional consequences of splicing dysregulation, we ran the MAJIQ 2.1 splicing algorithm on paired diagnostic/relapse samples while filtering for local splicing variations (LSV) with ≥20% difference in junction inclusion with 95% confidence (Supplementary Figure 1b). We further filtered for samples without known driver mutations and for LSVs affecting these wild-type driver genes. After performing hierarchical clustering, we discovered a cluster of 5 relapses with concurrent mis-splicing of multiple chemoresistance genes (Supplementary Figure 1c, light blue rectangle). To elucidate the underlying molecular mechanisms, we performed gene set enrichment analysis (18) on all transcripts differentially expressed in D-R pairs in that cluster of patients. Top 3 and 7 out of top 12 categories pertained to RNA splicing (Figure 1c). To determine whether AS is only in play in samples with unmutated drivers of chemoresistance or has a broader impact, we extended MAJIQ splicing analysis to all 48 D-R pairs. Unsupervised clustering re-discovered the same 5 patient samples and assigned additional 12 r/r leukemias to the same cluster for the total of 17 (Figure 1d, light blue rectangle), potentially implicating AS in ∼1/3 of B-ALL relapses. We obtained further evidence of SF dysregulation at the level of poison exon inclusion known to result in nonsense-mediated decay (NMD) (19). For example, in all relapse samples, almost 100% of transcripts encoding the aforementioned SRSF10 carried stop-codon-containing exon 3a, presumably rendering them non-functional (Supplementary Figure 1d).

We then considered more carefully the nature of AS events contributing to this cluster (Table 1). We noted that many of them mapped to genes conferring resistance to commonly used B-ALL therapeutics: glucocorticoids (*NR3C1*), thiopurines (*PRPS1 & NT5C2)*, and anti-folates (*FPGS*). In additional, they affected relapse genes associated with more general, not exclusively cancer cell-intrinsic mechanisms. Examples included the master epigenetic regulator *CREBBP* and its functional interactor *SETD2* and the universal pan-cancer suppressor *TP53*. Overall, the majority of the splicing variants in Table 1 are putative loss-of-function events, as they either directly disrupt open reading frames (mimicking nonsense mutations) or render the transcript vulnerable to NMD. For instance, *TP53* exon 9i contains a premature stop codon, and its inclusion results in the previously validated p53β protein isoform with multiple functional defects (20,21). However, the effects of *CREBBP* exon 25/26 skipping and the alternative 5’ splice site in *FPGS* exon 8 [previously reported in the literature (22,23)] could not be inferred *a priori* in the context of the loss-of-function model and required experimental investigation.

**Table 1.**
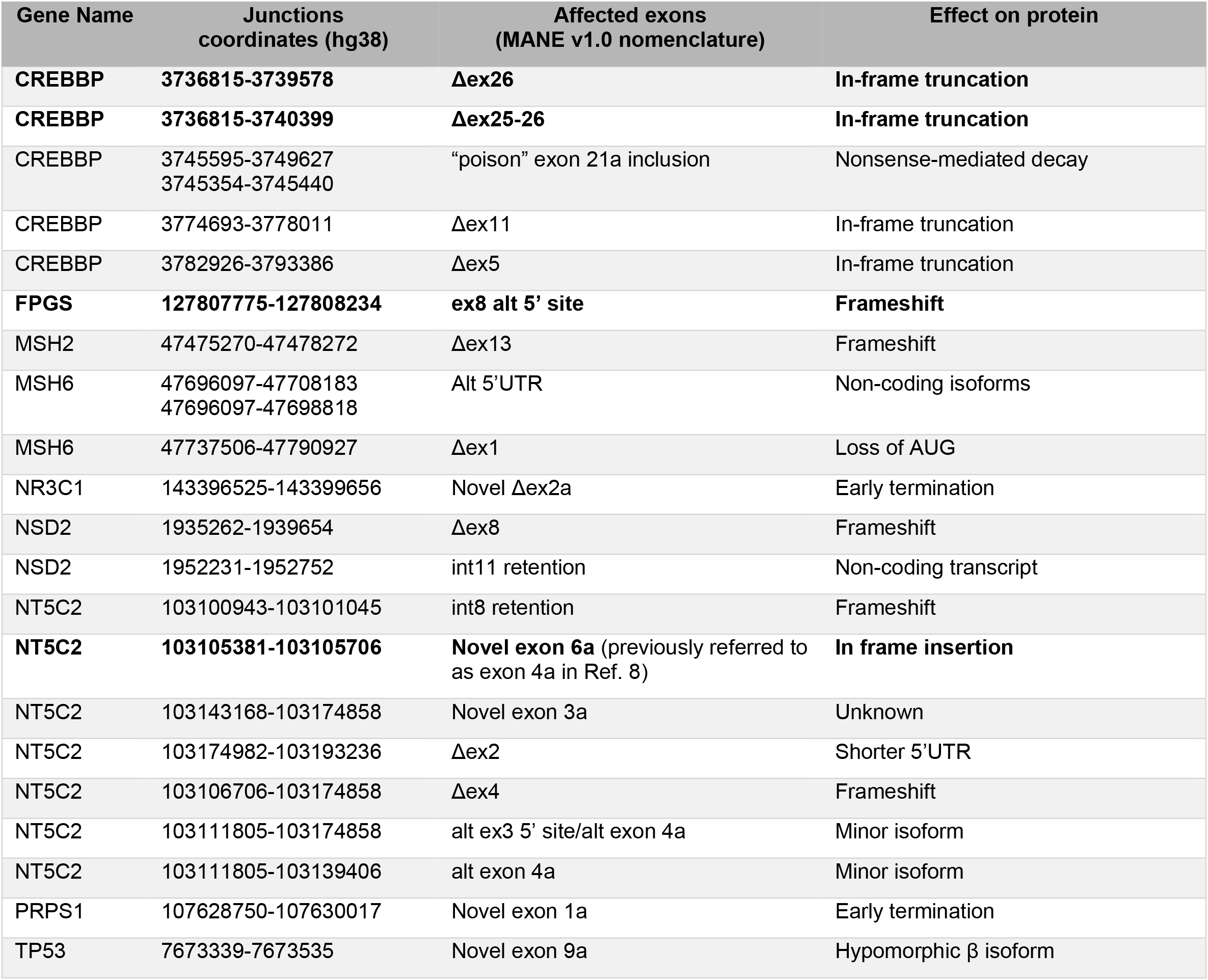
r/r B-ALL drivers affected by AS. Events **in bold** are under investigation in this study.

| Gene Name | Junctions coordinates (hg38) | Affected exons (MANE v1.0 nomenclature) | Effect on protein |
| --- | --- | --- | --- |
| **CREBBP** | **3736815-3739578** | **Δex26** | **In-frame truncation** |
| **CREBBP** | **3736815-3740399** | **Δex25-26** | **In-frame truncation** |
| CREBBP | 3745595-3749627<br>3745354-3745440 | “poison” exon 21a inclusion | Nonsense-mediated decay |
| CREBBP | 3774693-3778011 | Δex11 | In-frame truncation |
| CREBBP | 3782926-3793386 | Δex5 | In-frame truncation |
| **FPGS** | **127807775-127808234** | **ex8 alt 5' site** | **Frameshift** |
| MSH2 | 47475270-47478272 | Δex13 | Frameshift |
| MSH6 | 47696097-47708183<br>47696097-47698818 | Alt 5'UTR | Non-coding isoforms |
| MSH6 | 47737506-47790927 | Δex1 | Loss of AUG |
| NR3C1 | 143396525-143399656 | Novel Δex2a | Early termination |
| NSD2 | 1935262-1939654 | Δex8 | Frameshift |
| NSD2 | 1952231-1952752 | int11 retention | Non-coding transcript |
| NT5C2 | 103100943-103101045 | int8 retention | Frameshift |
| **NT5C2** | **103105381-103105706** | **Novel exon 6a** (previously referred to as exon 4a in Ref. 8) | **In frame insertion** |
| NT5C2 | 103143168-103174858 | Novel exon 3a | Unknown |
| NT5C2 | 103174982-103193236 | Δex2 | Shorter 5'UTR |
| NT5C2 | 103106706-103174858 | Δex4 | Frameshift |
| NT5C2 | 103111805-103174858 | alt ex3 5' site/alt exon 4a | Minor isoform |
| NT5C2 | 103111805-103139406 | alt exon 4a | Minor isoform |
| PRPS1 | 107628750-107630017 | Novel exon 1a | Early termination |
| TP53 | 7673339-7673535 | Novel exon 9a | Hypomorphic $\beta$ isoform |

### AS events of *CREBBP* and *FPGS* in r/r B-ALL are loss-of-function

While *CREBBP* mRNA was affected by several AS events, in-frame skipping of exons 25 and 26 was of potential functional significance, as it removes 261bp (87aa) from the HAT domain. This AS event was also among the most robust in our analysis of B-ALL relapses: on average, 50% of all *CREBBP* reads corresponded to the resultant ex24-ex27 junction (Figure 2a). In the representative example in Figure 2b, fewer than 30% of reads mapped uniquely to the Δex25/26 isoform in the diagnostic sample (32 reads in blue sashimi plots), whereas in the paired relapse sample this event predominated (99 reads in the red sashimi plots). To determine the effects of this event on protein function, we designed two sets of short guide RNAs mapping to the exons in question and used them to remove the exon 25-26 segment from 697 B-ALL cells via CRISPR-mediated genome editing. Using semi-quantitative RT-PCR, we detected decreased expression of the *CREBBP* full-length isoform and robust skipping of its exons 25 and/or 26, with combined skipping of both exons being the most readily detectable event (Figure 2c). This was also apparent when protein levels were analyzed using immunoblotting (Figure 2d, top panel). Since the splicing event affects the HAT domain, we chose, as a functional readout, its effects on histone modifications. We observed that *CREBBP* exon 25/26 skipping resulted in a commensurate reduction in H3K27Ac while preserving total H3 levels (Figure 2d). These results support the notion that *CREBBP* exon 25/26 skipping yields a protein isoform deficient in histone acetyl transferase activity.

**Figure 2.**
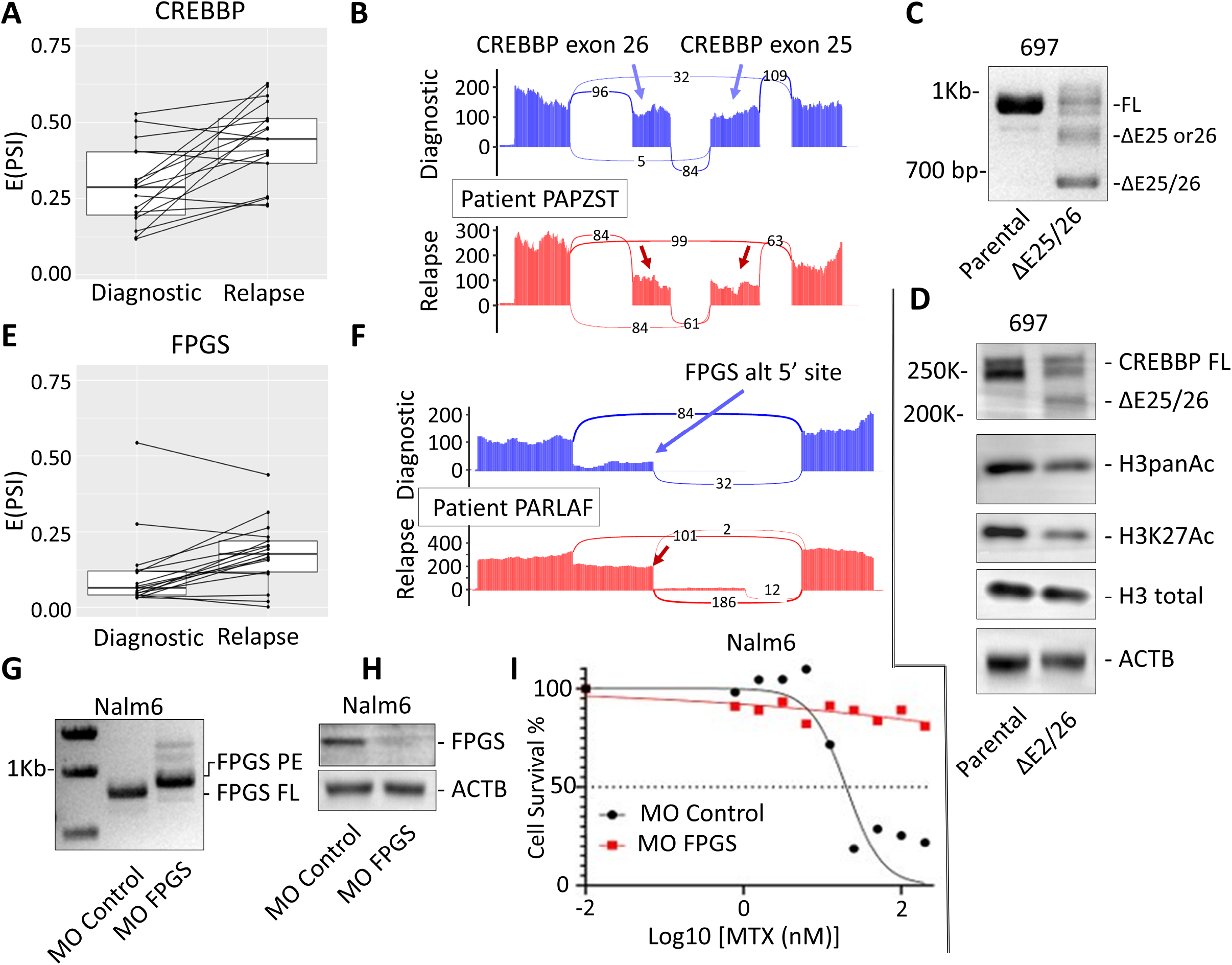
LOF AS events affecting CREBBP and FPGS. **a**. Box plot showing percentages of reads connecting CREBBP exon 24 to exon 27 in diagnosis/relapse pairs from panel 1d. “PSI” refers to percent-spliced-in. **b**, Sashimi plots visualizing the same CREBBP events in a representative PAPZST sample. **c**. RT-PCR analysis of CREBBP transcripts in 697 B-ALL cells transfected with exon 25 and 26-specific sgRNAs packaged into Cas9 particles. **d**. Immunoblotting analysis of the same samples using antibodies recognizing indicated proteins and post-translational modifications of histone H3. Beta actin was used as a loading control. **e**. Box plot showing percentages of reads connecting 5’ splice sites in FPGS exon 8 to the downstream exon 9 in diagnosis/relapse pairs from panel 1d. “PSI” refers to percent-spliced-in. **f**, Sashimi plots visualizing the same FPGS event in a representative PARLAF sample. **g**. RT-PCR analysis of FPGS transcripts in Nalm6 cells transfected with control or FPGS exon 8 5’ splice site-specific Morpholino. “FL” and “PE” refer to the full-length and the poison exon-containing isoforms, respectively. **h**. Immunoblotting analysis of the same samples using an anti-FPGS-antibody. **i**. IC50 plot representing survival of these cells after exposure to increasing concentrations of methotrexate (MTX).

The AS event affecting *FPGS* exon 8 was equally pervasive: only in a few pairs of samples was its direction reversed (Figure 2e). In a representative relapse sample depicted in Figure 2f, 65% of reads connected to the downstream exon 9 (186 in the red sashimi plot) originated at the alternative 5’ splice site, whereas in the paired diagnostic sample this ratio was reversed (32 reads in the blue sashimi plot). To model this event *in vitro*, we designed a morpholino (MO) antisense splice blocker targeting the canonical 5’ splice site, thus forcing the use of the alternative downstream site (see Materials and Methods). In transfected Nalm6 B-ALL cells, this MO almost completely redirected splicing towards the expected downstream site, resulting in partial intron inclusion and rendering exon 8 “poisonous” via inclusion of a stop codon (“PE” in Figure 2g). Therefore, the use of *FPGS* MO also resulted in a sharp decrease in FPGS protein levels, as evidenced by immunoblotting (Figure 2h). As FPGS is a known driver of sensitivity to methotrexate (24), we tested the effects of MO-induced FPGS AS on cell survival in response to this antifolate drug. In these experiments, redirecting splicing of *FPGS* exon 8 towards the downstream splice site resulted in essentially complete resistance to methotrexate and unattainable IC_50_ values (Figure 2i). These data solidified our conclusion that AS phenocopies loss-of-function mutations in this gene previously linked to chemoresistance in r/r B-ALL (5).

### AS of *NT5C2* yields an mRNA isoform conferring resistance to thiopurines

Across all relapse-associated AS events identified in our analysis (Table 1), the alternative isoform of *NT5C2* with the functionally uncharacterized in-frame exon 6a [previously referred to by us as exon 4a (8)] stood out. This microexon is annotated in RefSeq, and its robust inclusion was detected in normal bone marrow B cell progenitors in our earlier AS study (8) and in some other populations of hematopoietic cells in our recently unveiled MAJIQlopedia, a reference database for splicing variations across human cancers and normal tissues (25). However, it is not included in The Matched Annotation from the NCBI and EMBL-EBI (MANE), which is a broadly used standard for clinical reporting. Of note, increased frequencies of this event were observed in most relapsed B-ALL samples compared with the matched diagnostic specimens (Figure 3a). In a representative relapse sample depicted in Figure 3b, >80% of reads originating in the upstream canonical exon 6 (352 in the red sashimi plot) connected to exon 6a, whereas in the paired diagnostic sample <25% did (80 in the blue sashimi plot). This finding was consistent with our prior data showing lower levels of NT5C2 exon 6a inclusion in *de novo* leukemias compared to their normal bone marrow progenitors (8).

**Figure 3.**
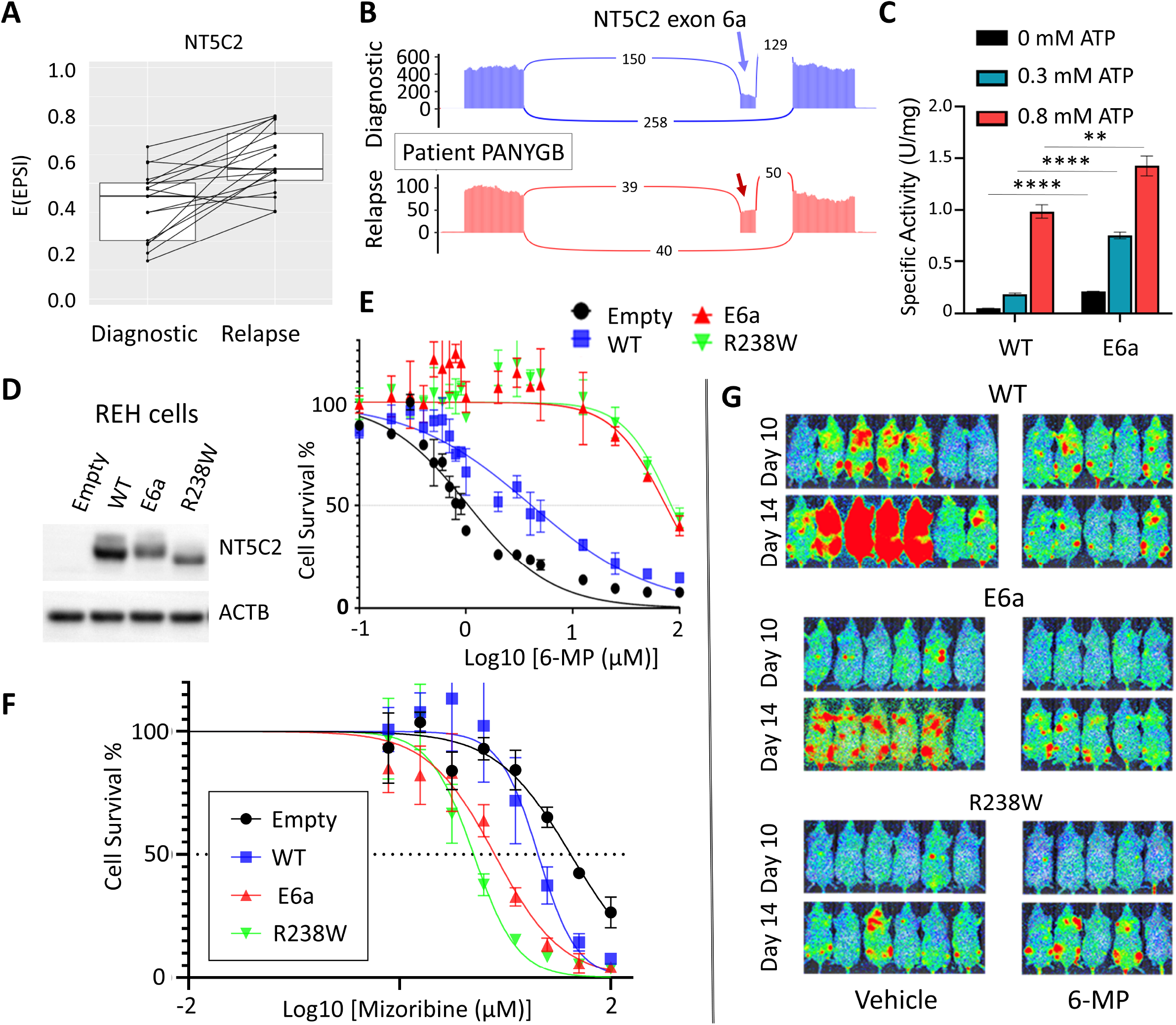
The NT5C2ex6a isoform as a driver of resistance to thiopurines. **a**. Box plot showing percentage of read connecting NT5C2 exon 6 to exon 6a in 6 diagnosis/relapse pairs depicted in panel 1c. “PSI” refers to percent-spliced-in. **b**. Sashimi plots visualizing the same NT5C2 events in a representative PANYGB sample. **c**. In vitro nucleotidase assays assessing the enzymatic activity of the canonical (WT) and E6a NT5C2 isoforms in the presence of increasing concentrations of ATP. Data are shown as mean ± SD. Asterisks indicate statistical significance per Student’s t-test. **d**. Expression levels of transduced NT5C2 isoforms in REH cells. “WT”, “E6a”, and “R238W” denote, respectively, the canonical isoform, the NT5ex6a splice isoform, and the R238W hotspot mutant. “Empty” denotes vector-only cells. **e**. The IC_50_ plot representing survival of these cells exposed to increasing concentrations of 6-MP. **f**. The IC_50_ plot representing survival of these cells exposed to increasing concentrations of mizoribine. **g**. Bioluminescent detection of the same cells additionally expressing the firefly luciferase gene. Cells were xenografted into NSG mice and imaged on days 10 and 14.

To determine whether *NT5C2* exon 6a can be found in translatable cap-to-poly(A) protein-coding transcripts we amplified its cDNA and subjected the resultant 1.7-kb fragment (Supplementary Figure 2a) to targeted re-sequencing using the Oxford Nanopore Technologies approach (see Supplementary Methods). Using this assay, we readily detected exon *NT5C2* 6a inclusion in the majority of reads corresponding to the Philadelphia chromosome-like MHH-CALL4 cell line with the IGH::CRLF2 rearrangement (Supplementary Figure 2b). Further support for the translatability of this isoform comes from its inclusion in the Uniprot database (accession code A0A6Q8PHP0) and the inclusion of the exon 6a-encoded tryptic peptide SQVAVQKR in The Peptide Atlas (accession code PAp02559851).

Inclusion of exon 6a in the *NT5C2* open reading frame results in the addition of 8 extra amino acids near the ATP-binding effector site 2. To determine the effect of this alteration on NT5C2 function, we produced the corresponding recombinant protein (Supplementary Figure 2c,d) and tested its enzymatic activity in nucleotidase assays with IMP as a substrate under basal conditions and following allosteric activation with ATP (14). In these cell-free assays, bacterially produced NT5C2ex6a exhibited elevated enzymatic activity compared with canonical NT5C2 protein in response to low concentrations of ATP (Figure 3c). This biochemical response was consistent with a gain of function mechanism, similar to that observed for most relapsed-associated *NT5C2* mutations characterized to date (14).

To rigorously test a potential role of *NT5C2* ex6a in 6-MP resistance, we generated REH B-ALL cell models expressing canonical *NT5C2, NT5C2* ex6a, and *NT5C2* R238W, a validated gain-of-function resistance driver with the most common hotspot mutation (14) (Figure 3d). Cell viability assays carried out with increasing concentrations of 6-MP demonstrated that expression of the *NT5C2* ex6a isoform confers 6-MP resistance comparable to that induced by the *NT5C2* R238W mutation and well above that conferred by the canonical isoform of wild-type *NT5C2* (Figure 3e and Supplementary Figure 2e), although the effects were somewhat less potent in Nalm-6 cells (Supplementary Figure 2f). Metabolically, increased enzymatic activity of NT5C2 mutants results in increased clearance of thiopurine mononucleotide monophosphate metabolites, but also in depletion of intracellular nucleotide pools, which in turn renders leukemia cells more sensitive to mizoribine, an inosine monophosphate dehydrogenase (IMPDH) inhibitor (16). Therefore, we evaluated the relationship between *NT5C2* ex6a and sensitivity to this drug. Once again, both *NT5C2* R238W- and *NT5C2* ex6a-expressing cells showed convergent phenotypes with respect to increased sensitivity to mizoribine treatment *vis-a-vis* cells expressing the canonical isoform of wild-type *NT5C2* (Figure 3f and Supplementary Figure 2g).

We further noted that the NT5C2 exon 6a-encoded peptide SQVAVQKR contains a characteristic SQ motif, which constitutes the recognition site for the DNA-PK/ATR/ATM subfamily of kinases of the PIKK family (26), also reflected in the Kinase Library (27). This observation suggested a potential role for protein phosphorylation at this site in the regulation of NT5C2 activity. Thus, we generated mutant variants of NT5C2 ex6a, in which the ex6a-encoded serine residue was replaced by the common phosphomimetic aspartic acid (S131D) or by alanine (S131A), to serve as a phosphorylation-deficient control. Evaluation of responses to 6-MP treatment of REH cells expressing each of these constructs revealed that while the parental NT5C2 E6a isoform and its S131A derivative conferred equivalent levels of 6-MP resistance, the S131D substitution resulted in the markedly increased resistance phenotype, with one log further increase in IC50 (Supplementary Figure 2h). These results suggest that phosphorylation events at the S131 site can augment *NT5C2* exon 6a-driven 6-MP resistance.

To directly assess the impact of *NT5C2* ex6a inclusion on leukemia cells growth and therapy responses *in vivo*, we implanted luciferase-expressing REH cells expressing the canonical wild type *NT5C2* isoform, *NT5C2* R238R, and *NT5C2* ex6a into NSG mice and imaged animals on days 5, 8, 12, and 14. Previous reports linked hyperactive NT5C2 with a “fitness cost” phenotype and decreased cell proliferation resulting from the depletion of intracellular nucleotide pools (16). Consistent with these observations, expression of *NT5C2* R238W and *NT5C2* ex6a induced clear growth retardation when compared with expression of the canonical *NT5C2* isoform (Supplementary Figure 3a). Nevertheless, we observed robust leukemia engraftment, allowing us to test the effect of the three different *NT5C2* variants on chemotherapy response by treating each experimental group with either vehicle or 6-MP (Figure 3g). When compared to the initial treatment timepoint (day 10 post-engraftment), the WT model shows a negative fold change (plotted on the Y-axis in Supplementary Figure 3b), which was indictive of cell death. In contrast, the E6a model demonstrates slower cell division under treatment, but the fold change remains non-negative, signifying that the treatment does not induce cell death. Notably, the E6a model exhibits an identical fold change to the R238W mutant model. These findings further support the comparable roles of these two NT5C2 variants in promoting chemoresistance.

## Discussion

RNA-centric regulatory mechanisms, such as formation of fusion transcripts and circular RNAs, RNA editing, and in particular aberrant mRNA splicing, recently have been recognized as key drivers of neoplastic growth (28). Indeed, mutations in genes encoding splicing factors (SF3B1, SRSF2, etc.) are very common in adult leukemias, raising the possibility that targeting aberrant splicing programs could be a viable therapeutic option. Although such mutations are exceedingly rare in pediatric AML and ALL, a growing body of work by us and others has revealed that inclusion of “poison” and other non-canonical exons in SF-encoding transcripts is among the most consistent splicing aberration in both B- ALL and T-ALL (8,29) as well as in many solid tumors (19,30). Additionally, we show that in TARGET r/r B-ALL samples, SRSF- and HNRNP-encoding genes are frequently affected by copy number alterations such as deep deletions, likely exacerbating AS.

We also report pervasive dysregulation, at the level of mRNA splicing, of key chemosensitivity genes. In most cases (CREBBP, FPGS, and other examples in Table 1) the AS events are of the loss-of-function nature, making it challenging to devise pharmacological interventions. Still, an RNA-based diagnostic test detecting, for example, the heavy usage of FPGS ex8 alt 5’ site could guide clinical decisions regarding the use of methotrexate. Thus, we envision that targeted RNA-seq panels, either short- or long-read, will become valuable companion diagnostics, despite known limitations of underlying technologies (31).

The case of NT5C2 is particularly interesting from the therapeutic standpoint, since NT5C2 splicing aberrations leading to the inclusion of cryptic exon 6a ultimately create a proteoform with gain-of-function properties. Since gain-of-function NT5C2 variants interfere with cell proliferation (16), it comes as no surprise that the inclusion of NT5C2 exon 6a is actually reduced in *de novo* leukemias compared to pro-B cells (8) and is re-gained only upon treatment with thiopurines, as a trade-off between survival and proliferation. This complex dynamic was apparent in our in vivo experiments, where both NT5C2 E6a and R238W leukemias grew more slowly in NSG mice than the controls but effectively resisted treatment with 6-MP. Of interest, at least in one leukemia (PARJZZ) NT5C2 exon 6a inclusion co-exists with the hotspot mutation R367Q reported earlier (4). This finding is consistent with an evolutionary model wherein non-mutational mechanisms (alternative splicing, NT5C2 phosphorylation) confer initial resistance to 6-MP, enable persistence of proliferating cells, and allow eventual emergence of clones with gain-of-function NT5C2 mutations and a constitutive resistance phenotype.

Robust expression of the NT5C2ex6a isoform not only predicts resistance to thiopurines, but may suggest additional therapeutic options, such as treatment with the emerging class of direct NT5C2 small-molecule inhibitors such as CRCD2 (32) and also with clinically used IMPDH inhibitors and immunosuppressive agents mycophenolate mofetil (MMF) and mizoribine (33). In support of this notion, NT5C2ex6a-expressing REH and NALM6 cells were found to exhibit heightened sensitivity to mizoribine compared to the isogenic cell lines reconstituted with the canonical NT5C2 isoform.

Other ways to therapeutically target NT5C2 dysregulation are likely to exist. Our recently published work elucidated the role of NT5C2 post-transcriptional modification in chemoresistance, with phosphorylation of the Ser-502 residue by a yet-to-be-identified protein kinase increasing its activity against 6-MP (32). Our new data showing that substituting Ser at position 131 with a phosphomimetic Asp augments NT5C2ex6a-induced resistance to 6-MP lend further credence to the model linking NT5C2 phosphorylation and chemoresistance function. If further research confirms the involvement of ATR/ATM family kinases in phosphorylating the NT5C2 exon 6a-encoded SQ motif, a variety of small-molecule inhibitors of these enzymes (34) will be available for preclinical studies and clinical trials aiming to re-sensitize r/r B-ALL to chemotherapy and in doing so improve outcomes in this aggressive and often fatal childhood cancer.

## Acknowledgements

The authors are grateful to past and present members of the Barash, Tasian, and Thomas-Tikhonenko laboratories and Penn’s RNA Club for many helpful discussions. This work was supported by grants from the NIH | National Cancer Institute (U01 CA232563 to ATT and YB, U01 CA232486 and U01 CA243072 to SKT, and P30 CA013696, R35 CA210065, and CA206501 to AF), Pennsylvania Department of Health SFY22 CURE Non-Formula Collaborative Research on Childhood and Adolescent Blood Cancers (#67-173 to ATT), United States Department of Defense (CA180683P1 to SKT), St. Baldrick’s Foundation (EPICC Team and St. Baldrick’s-Stand Up to Cancer Dream Team Translational Cancer Research Grant [SU2C-AACR-DT-27-17] to ATT and SKT), The V Foundation for Cancer Research (T2018-014 to ATT), The Emerson Collective (886246066 to ATT), Alex’s Lemonade Stand Foundation (Innovation Awards to ATT and AF), and Leukemia & Lymphoma Society (Translational Research Grant 6455–15 and Screen to Lead Grant 8011–18 to AF). The indicated SU2C research grant is administered by the American Association for Cancer Research, the scientific partner of SU2C. Additionally, CR is a past Special Fellow and SKT is a Scholar of the Leukemia & Lymphoma Society. ATT and AF further acknowledge support from the SPROUT Program and Accelerating Cancer Therapeutics (ACT) Program at CHOP and Columbia University, respectively. SKT is Joshua Kahan Endowed Chair in Pediatric Leukemia Research and ATT is Mildred L. Roeckle Endowed Chair in Pathology at Children’s Hospital of Philadelphia.

