## Supplementary Methods and Data for "An Alternatively Spliced Gain-of-Function NT5C2 Isoform Contributes to Chemoresistance in Acute Lymphoblastic Leukemia"

#### Supplemental Materials and Methods

***RNA-seq analysis.*** RNA-seq reads were first trimmed to remove adapters (BBTools version 38.87) and were then aligned using STAR version 2.7.3a to the hg38 reference genome, while providing known gene isoforms through the GENCODE annotation version 32. Additionally, we used STAR flags “--quantMode GeneCounts” and “--alignSJoverhangMin 8” to quantify genes and ensure spliced reads had an overhang of at least 8 bases. Junction spanning reads were obtained from the STAR “*_SJ.out.tab” result files and each entry was normalized by dividing by the total number of junction-spanning reads and multiplying by a factor of one million, yielding a JPM value. Visualization and downstream analyses were conducted in R using the ggplot2 and tidyverse packages.

***Mutation analysis.*** Relapse-specific variant discovery was done using the Genome Analysis Toolkit (GATK) version 4.4.0.0 (RRID:SCR_001876) on the RNA-seq data. Shortly, aligned reads were marked for duplicated events using Picard Tools version 3.0.0. Reads spanning splicing events were split into independent reads with the SplitNCigarReads tool followed by base recalibration. To specifically call mutations that are enriched in the relapsed samples, the Mutect2 algorithm was applied using the corresponding diagnostic sample to each relapsed file in the –normal flag. All the variants were annotated using the Funcotator tool with the hg38 somatic annotation from the FuncotatorDataSourceDownloader. All the maf files from the 48 patients were loaded into the R package maftools for variant visualization and downstream analysis.

***qRT-PCR.*** Total RNA was isolated using Maxwell RSC simplyRNA Cells kit with the Maxwell RSC48 Instrument (Promega) and reverse-transcribed using SuperScript IV (Invitrogen; 18090010). Primers used for mRNA isoform quantification are listed in Supplementary Tables. qRT-PCR was performed using PowerSYBR Green PCR Master Mix (Life Technologies). Reactions were performed on an Applied Biosystems Viia7 machine and analyzed with Viia7 RUO software (Life Technologies). When indicated, individual PCR products were gel-purified (QIAquick Gel Extraction Kit; Qiagen) and Nanopore sequenced.

***Western blotting.*** For the Western blotting, cells were lysed in RIPA buffer plus protease and phospatase inhibitor and benzonase. 30ug of protein were loaded on NuPAGE gels. For detection by Western blotting, the following primary antibodies were used: anti-NT5C2 antibody (Sigma-Aldrich Cat# WH0022978M2, RRID:AB_1842747), anti-FPGS antibody (Abcam Cat# ab184564 [clone EPR11064], RRID:unknown), anti-CREBBP (Cell Signaling Technology Cat# 7389, RRID:AB_2616020), anti-histone H3 acetyl K9 + K14 + K18 + K23 + K27 (Abcam Cat# ab47915, RRID:AB_873860), anti-histone H3 acetyl K27 (Cell Signaling Technology Cat# 8173, RRID:AB_10949503), and anti-histone H3 (Active Motif Cat# 61475, RRID:AB_2687473) in combination with anti-rabbit or anti-mouse HRP-linked secondary antibodies (Cell Signaling Technology) and Amersham’s enhanced chemiluminescence Western blotting detection reagent (GE Life Sciences). Additionally, a horseradish peroxidase-conjugated antibody was used to detect the β-actin loading control (Cell Signaling Technology Cat# 12262, RRID:AB_2566811).

##### *PCR and Sequencing Primers*

| Primer_Name | Sequence |
|---|---|
| GAPDH_qPCR_F | ATGGGGAAGGTGAAGGTCG |
| GAPDH_qPCR_R | GGGGTCATTGATGGCAACAATA |
| ACTB_qPCR_F | AGCATCCCCCAAAGTTCAC |
| ACTB_qPCR_R | AAGGGACTTCCTGTAACAACG |
| NT5C2_qPCR_E4E4a_F1 | TTCCCTACCAGGGGACTTGT |
| NT5C2_qPCR_E4E4a_R1 | GAACAGCTACCTGAGAACCCC |
| NT5C2_qPCR_E4E4a_F2 | ACATTCCCTACCAGGGGACT |
| NT5C2_qPCR_E4E4a_R2 | ACAGCTACCTGAGAACCCCTT |
| NT5C2_qPCR_TOTAL_F1 | CGCACAGGAGCTACATGTCT |
| NT5C2_qPCR_TOTAL_R1 | TGGAACTGATGTCTGGACGC |
| NT5C2_qPCR_TOTAL_F2 | CAAAGAAACGGCAAGGGTGG |
| NT5C2_qPCR_TOTAL_R2 | TGCTTGTAGAGTTCAGCCAAGA |
| NT5C2_qPCR_E4E5_F1 | AACTTTATAAGGGGACCAGAAACT |
| NT5C2_qPCR_E4E5_R1 | CTAGGCAGGCCAACAGGTAG |
| NT5C2_qPCR_E4E5_F2 | GGGGACCAGAAACTAGAGAACA |
| NT5C2_qPCR_E4E5_R2 | CTAGGCAGGCCAACAGGTAG |
| NT5C2_cloning_F | ATGTCAACCTCCTGGAGTGATC |
| NT5C2_cloning_R | TTGTTCCTGTGAGTCCTGCC |
| NT5C2_E4a_cloning_F | TTCTCAGGTAGCTGTTCAGAAGAGACCAGAAACTAGAGAACAGTATCCAA |
| NT5C2_E4a_cloning_R | CTCTTCTGAACAGCTACCTGAGAACCCCTTATAAAGTTAAATCCATGTGC |
| NT5C2_R238W_cloning_F | GCTTCTGAGCtGGATGAAGGAAGTAGGGAAAGTATTTCTT |
| NT5C2_R238W_cloning_R | CCTTCATCCaGCTCAGAAGCAAAGGCAGTTTTCCATCTTT |
| FPGS_NANOPORE_F | GGGGCTTATGACTGCACCAA |
| FPGS_NANOPORE_R | CTTCGTCCAGGTGGTTCCAG |

##### *sgRNA for Genome Editing*

| sgRNA_Name | Sequence |
|---|---|
| sgRNA_Exon26_downstream | GTTTACCATGAGATCCTTAT |
| sgRNA_Exon26_upstream | AGAAATGTACACACGCCTGT |
| sgRNA_Exon25_downstream | AACTACCTCGTGTTTGGAGG |
| sgRNA_Exon25_upstream | CCAGAGCCGGACATTTACAA |

### Supplemental Figures

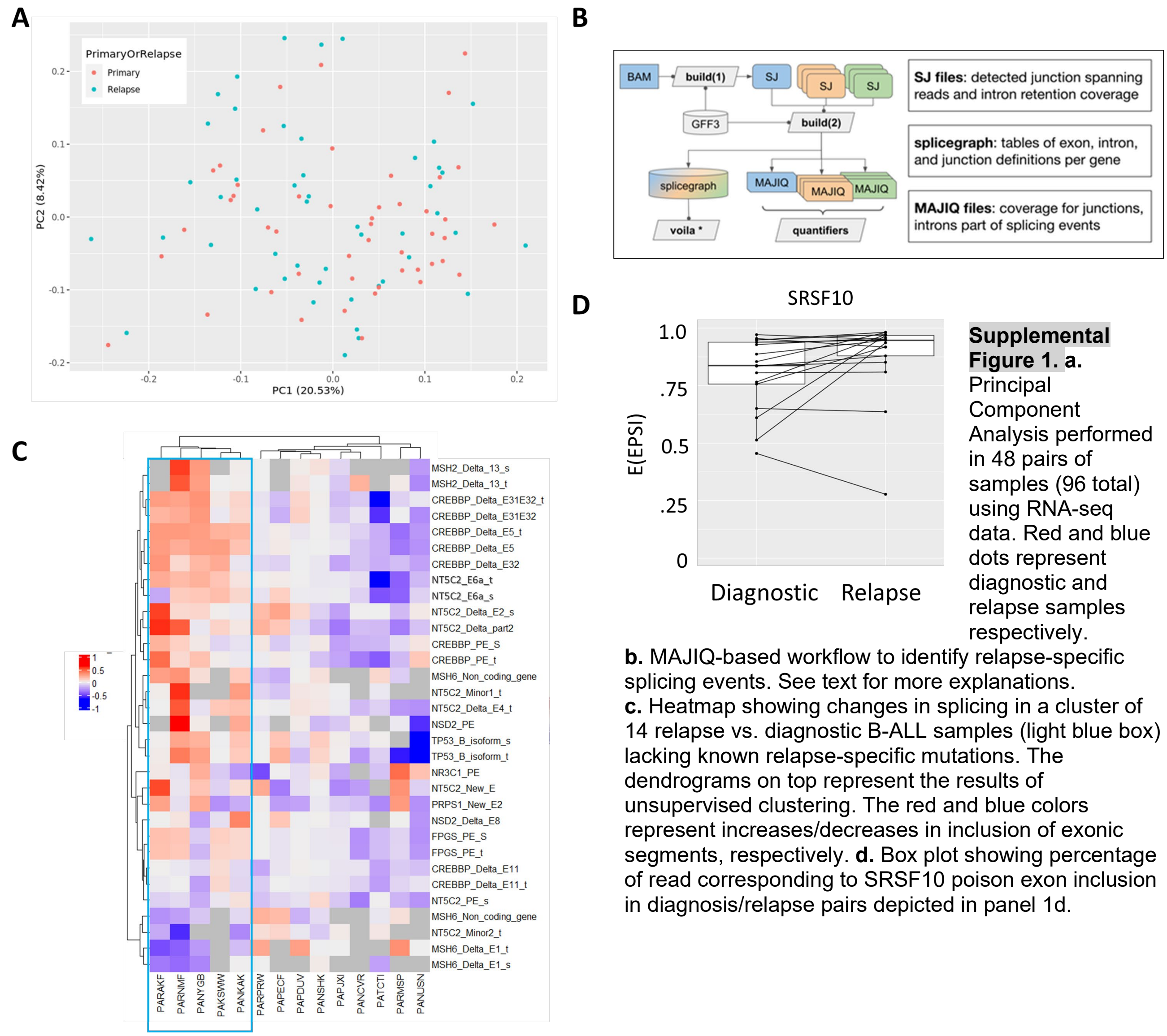


**Supplemental Figure 1. a.** Principal Component Analysis performed in 48 pairs of samples (96 total) using RNA-seq data. Red and blue dots represent diagnostic and relapse samples respectively.
**b.** MAJIQ-based workflow to identify relapse-specific splicing events. See text for more explanations.
**c.** Heatmap showing changes in splicing in a cluster of 14 relapse vs. diagnostic B-ALL samples (light blue box) lacking known relapse-specific mutations. The dendrograms on top represent the results of unsupervised clustering. The red and blue colors represent increases/decreases in inclusion of exonic segments, respectively. **d.** Box plot showing percentage of read corresponding to SRSF10 poison exon inclusion in diagnosis/relapse pairs depicted in panel 1d.

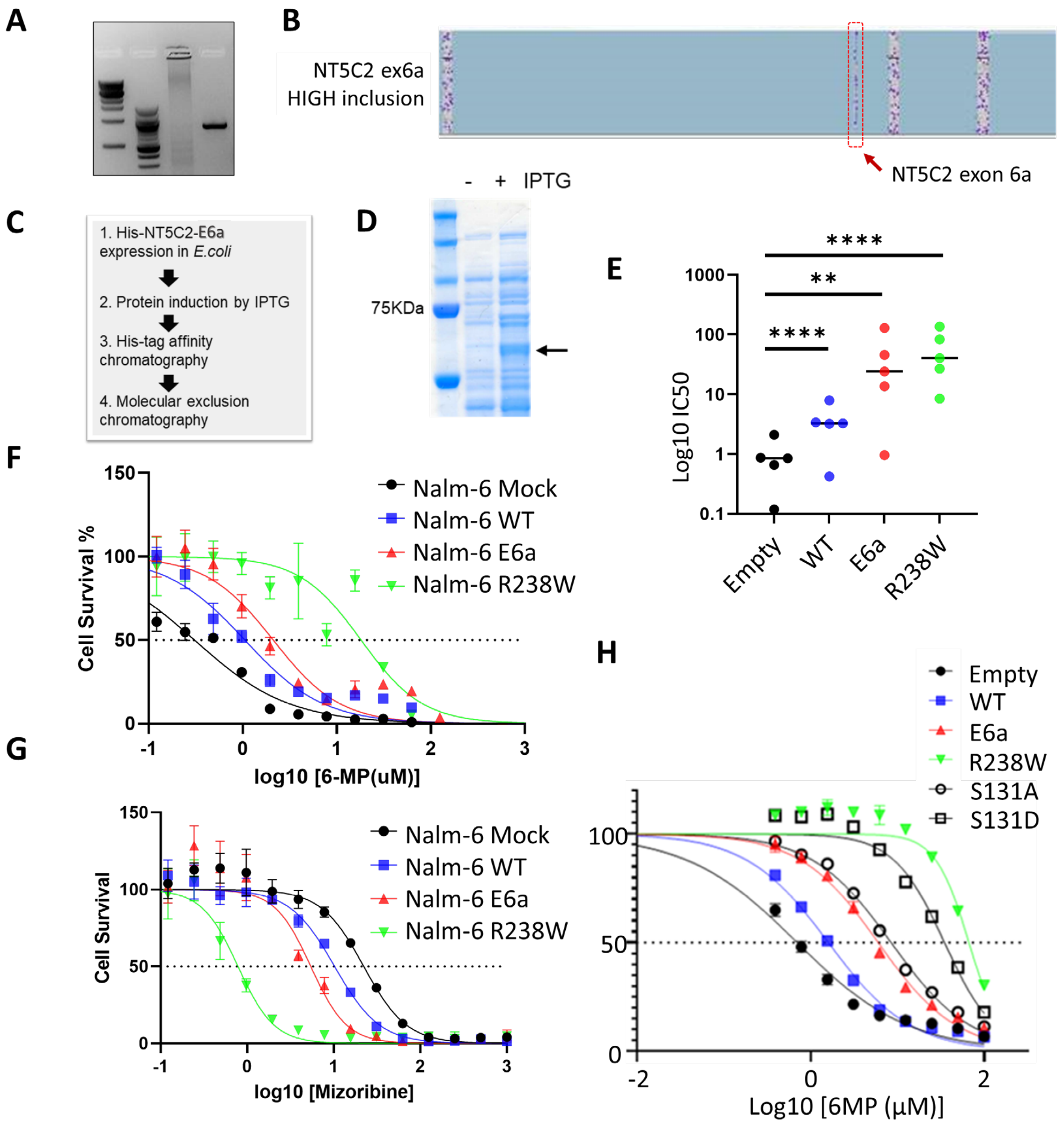


**Supplemental Figure 2. The role of NT5C2ex6a in chemoresistance in vitro.** **a.** RT-PCR amplification of the full-length NT5C2ex6a cDNA from MHH-CALL4 cells. **b**. Targeted long-read ONT sequencing of the NT5C2ex6a. **c.** General schema of NT5C2 ex6a protein purification. **d.** Electrophoretic detection of the NT5C2 ex6a isoform (arrow) following induction by IPTG. **e.** Average IC50 values from 5 independent experiments depicted in Figure 3e. Asterisks indicate statistical significance per Student's t-test. **f.** The $IC_{50}$ plot representing survival of NT5C2-expressing Nalm6 cells exposed to increasing concentrations of 6-MP. **g.** The $IC_{50}$ plot representing survival of NT5C2-expressing Nalm6 cells exposed to increasing concentrations of mizoribine. **h.** The $IC_{50}$ plot representing survival of additional NT5C2 variant-expressing REH cells exposed to increasing concentrations of 6-MP.

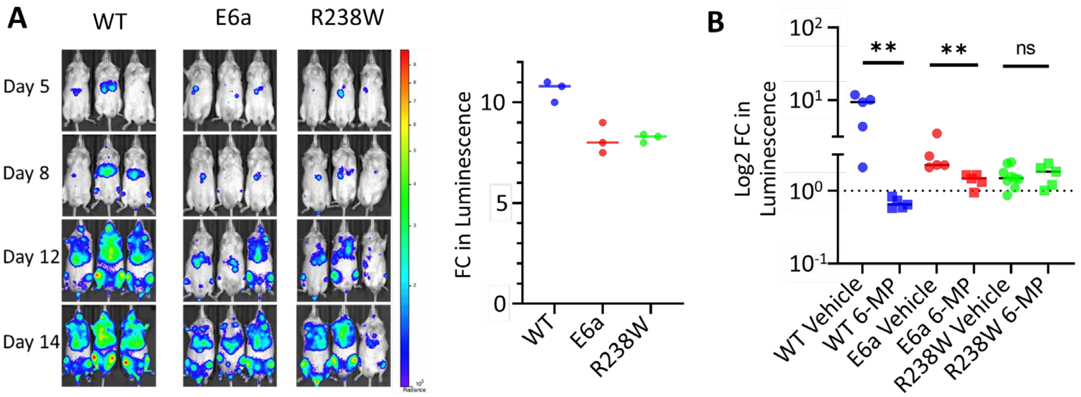


**Supplemental Figure 3. The role of NT5C2ex6a in chemoresistance in vivo.** **a**. LEFT: Ivis bioluminescence image of mice engrafted with REH Luc NT5C2 WT, E6a or R238W cells at the indicated time points. RIGHT: Quantitation of data from the previous panel, expressed as fold change in photon counts between d14 and d5. **b.** Quantitation of data from Figure 3g, expressed as fold change in photon counts between d14 and d10. Asterisks indicate statistical significance per Student's t-test.
